# Task-induced deactivation in diverse brain systems correlates with interindividual differences in distinct autonomic indices

**DOI:** 10.1101/275826

**Authors:** Vittorio Iacovella, Luca Faes, Uri Hasson

**Affiliations:** Center for Mind/Brain Sciences, The University of Trento, Trento, Italy; BIOtech, Department of Industrial Engineering, University of Trento, Trento, Italy; IRCS PAT-FBK Trento, Italy; Center for Practical Wisdom, The University of Chicago, Chicago (USA)

**Keywords:** ANS, Deactivation, Arithmetic, Interindividual Differences

## Abstract

Neuroimaging research has shown that different cognitive tasks induce relatively specific activation patterns, as well as less task-specific deactivation patterns. Here we examined whether individual differences in Autonomic Nervous System (ANS) activity during task performance correlate with the magnitude of task-induced deactivation. In an fMRI study, participants performed a continuous mental arithmetic task in a task/rest block design, while undergoing combined fMRI and heart / respiration rate acquisitions using photoplethysmograph and respiration belt. As expected, task performance increased heart-rate and reduced the RMSSD, a cardiac index related to vagal tone. Across participants, higher heart rate during task was linked to increased activation in fronto-parietal regions, as well as to stronger deactivation in ventromedial prefrontal regions. Respiration frequency during task was associated with similar patterns, but in different regions than those identified for heart-rate. Finally, in a large set of regions, almost exclusively limited to the Default Mode Network, lower RMSSD was associated with greater deactivation, and furthermore, the vast majority of these regions were task-deactivated at the group level. Together, our findings show that inter-individual differences in ANS activity are strongly linked to task-induced deactivation. Importantly, our findings suggest that deactivation is a multifaceted construct potentially linked to ANS control, because distinct ANS measures correlate with deactivation in different regions. We discuss the implications for current theories of cortical control of the ANS and for accounts of deactivation, with particular reference to studies documenting a “failure to deactivate” in multiple clinical states.

## 1 Introduction

Understanding the computations that different brain regions perform during task performance continues being one of the main undertakings of modern cognitive neuroscience. In investigating this issue, multiple studies show that task-evoked changes are not limited to brain networks strictly associated with task-related information processing. Specifically, engagement in different tasks (e.g., language, attention, memory) impacts activity and connectivity (e.g., Fransson & Marrelec, 2008; McKiernan, D’Angelo, Kaufman, & Binder, 2006; McKiernan, Kaufman, Kucera-Thompson, & Binder, 2003) of a “default mode” network (DMN) that is thought to mediate spontaneous, task-independent computations such as mind wandering (Mason et al., 2007).

Another way by which tasks can perturb brain activity is by modulating brain networks involved in Autonomic Nervous System (ANS) monitoring and regulation, which are often found to be distinct from the DMN in terms of topology and function (as we review below). The ANS is strongly impacted by tasks that present arousing stimuli or that specifically employ emotional stressors (e.g., Thayer, Ahs, Fredrikson, Sollers, & Wager, 2012; Thayer, Hansen, Saus-Rose, & Johnsen, 2009). However, the ANS is also perturbed by affectively neutral tasks such as mental arithmetic or Stroop tasks (for review, see Beissner, Meissner, Bar, & Napadow, 2013). Such perturbations have been linked to fluctuations in task performance (see Critchley & Garfinkel,2015, for review).

Our general aim in the current study was to establish how individual-differences in ANS activity relate, if at all, to BOLD deactivation during an affectively neutral task. Specifically, we examined whether there are brain networks where the magnitude of task-induced deactivation is associated with inter-individual differences (IID) in ANS reactivity during performance of a simple continuous mental arithmetic task. Our study addresses two fundamental issues concerning the relation between IID in ANS activity and task-related deactivation (as well as task-related activation). First, as we review below, few prior studies have specifically taken on a systematic examination of whether IID in ANS activity are related to task-induced activation or deactivation. And second, those studies that examined this question had relied on a single autonomic measure. Consequently, whether the magnitude of task-related deactivation in different brain systems is associated with different ANS indices, is a topic simply not addressed to date.

Interestingly, there exist marked IID in both ANS reactivity and task-induced deactivation, suggesting these may load on shared factors. Inter-individual differences in ANS reactivity are not only prevalent within non-clinical participant groups (Goldberger, Challapalli, Tung, Parker, & Kadish, 2001; Karemaker & Wesseling, 2008), but also vary with age (Pfeifer et al., 1983), and with personality features during development (Beauchaine, 2001). Altered ANS function is also associated with clinical states such as autism (Hirstein, Iversen, & Ramachandran, 2001) or depression (Carney, Freedland, & Veith, 2005). In tandem, IID in task-related (de)activation have also been reported, and these have been associated with similar factors to those that impact ANS. For instance, IID in activation/deactivation have been linked to stress level (Soares et al., 2013), age (Persson, Lustig, Nelson, & Reuter-Lorenz, 2007) meditation (Lutz, Brefczynski-Lewis, Johnstone, & Davidson, 2008) or clinical states such as autism and schizophrenia that have been linked to a “failure to deactivate” during simple cognitive tasks (e.g., Kennedy, Redcay, & Courchesne, 2006; Landin-Romero et al., 2015). Interestingly, IID in task-related activation/deactivation also correlate with differences in resting-state (baseline) fluctuation levels (Zou et al., 2013), which in turn are also linked to IID in ANS activity (Jennings, Sheu, Kuan, Manuck, & Gianaros, 2016). We therefore hypothesized that IID in ANS reactivity, as measured in a normal non-clinical group of participants, could be related to the extent of task-induced activation/deactivation.

Several studies have reviewed the brain systems involved in regulation of the ANS, particularly from the perspective of the psychology of emotion, or the involvement of ANS in interpersonal interactions (for reviews and meta-analyses, see, Beissner, Meissner, Bar, & Napadow, 2013; Thayer, Ahs, Fredrikson, Sollers, & Wager, 2012). The relation between ANS function and brain activity during simple cognitive tasks has received less investigation. Beissner et al. (2013) reported a meta-analysis of neuroimaging studies that used motor, emotional or cognitive stressors. The meta-analysis showed that affective, motor and cognitive tasks produce different relations between ANS activity and brain activity (see also Thayer et al., 2012, for a meta-analysis focusing on parasympathetic correlates). Particularly relevant for our current inquiry, Beissner et al. showed that during cognitive-task stressors, parasympathetic responses were linked to the left amygdala and right anterior insula. Sympathetic activity was linked to mid-cingulate cortex, left anterior insula, left secondary somatosensory cortex, vmPFC, subgenual ACC, left superior parietal lobule, left surpramarginal gyrus, and left amygdala.

We note that the relation between IID in ANS responses and task-related activation has been largely ignored in many prior neuroimaging studies, specifically because typical analyses collapse across these differences. As such, the bulk of prior work has focused on identifying brain regions where activity covaries with ANS fluctuations at the group level, and so between-participant differences were modeled as random effects. This was achieved, for example, by including time-series of ANS fluctuations as an explanatory variable (regressor) in single-subject BOLD-fMRI regression models (Critchley et al., 2003; Critchley, Tang, Glaser, Butterworth, & Dolan, 2005; Evans et al., 2009; Napadow et al., 2008), using block-mean ANS measures as parametric modulators in block designs (Fechir et al., 2010) or establishing ANS/PET-rCBF correlations on the single-participant level (Gianaros, Van Der Veen, & Jennings, 2004; Lane et al., 2009). Crucially, in all these analyses, the single-participant regression coefficients were used in second-level analyses to identify ANS-related activity at the group level, and IID were not quantified.

Furthermore, some neuroimaging studies that did report BOLD correlates of IID in ANS, limited their analysis to brain areas strongly implicated in the experimental task studied. To illustrate, Matthews et al. (2004) identified clusters sensitive to congruence in a Stroop task, and only within those did they evaluate correlations between heart-rate variability (HRV) and response levels to congruent and incongruent trials. This approach may be less sensitive to identifying activity-correlates of IID in ANS, as brain areas that are most strongly task-activated, or most strongly discriminative of two conditions at the group level may be those least impacted by arousal. In another fMRI study (Muehlhan et al., 2013) the authors examined IID in sympathetic responses as measured by salivary alpha amylase (sAA) while participants responded to validly- or invalidly-cued targets. The authors identified brain regions that satisfied two criteria: task-induced activity changes (either activation or deactivation) and significant BOLD/sAA correlations. Perhaps due to the motor-component of this task, the task-active regions that also showed correlations with sAA were not ones typically associated with the ANS. However, the authors identified several task-deactive regions including the left precuneus, angular gyrus bilaterally, vmPFC and left middle frontal gyrus. In all these regions, higher arousal was associated with greater deactivation. This and prior work (Nagai, Critchley, Featherstone, Trimble, & Dolan, 2004; Wong, Mass, Kimmerly, Menon, & Shoemaker, 2007) suggests that increased arousal is associated with greater disturbance of the ‘default’ process mediated by these regions.

To our knowledge, two studies have specifically treated the issue of IID in ANS as the focus of investigation. Wager et al. (2009) found that during responses to social threat, rostral and pregenual ACC showed a positive correlation between heart rate reactivity and task-induced activation. The right orbitofrontal cortex was deactivated by the task and showed an inverse correlation, so that heart rate reactivity was associated with greater deactivation. Importantly, in both regions, rapid BOLD fluctuations tracked fluctuations in heart rate, indicating that deactivated regions may track ANS fluctuations on a fine temporal scale. Gianaros et al. (2012)examined IID in beat-to-beat blood pressure and interbeat intervals (measurements obtained in mock fMRI scanner). They correlated a mental effort index of the task (effect size for *di*ffi*cult task easy task*) with IID in ANS reactivity. Across participants, stronger effect sizes were associated with stronger suppression of baroreflex sensitivity [BRS], an ANS index reflecting the short-term homeostatic control of blood pressure. This was found across the cingulate cortex but also in the insula and midbrain. The study considered however a single physiological index (BRS), which is also difficult to measure within a scanning environment thus reducing its applicability for everyday neuroimaging studies.

To summarize, there is limited understanding of how task-induced deactivation is related to IID in different autonomic measures. This holds particularly for emotionally-neutral cognitive tasks and for ANS-related metrics that can be established from typical cardiac and respiratory in-scanner recordings. To examine this issue, we established the magnitude of task-induced activation (or deactivation), and evaluated it against several ANS measures. We selected measures that differentially load on sympathetic and parasympathetic sources, and that can be derived from physiological signals collected concurrently with task performance in an fMRI scanner. Importantly, because we wanted to know whether IID correlations between ANS and task-induced effects are mainly found in areas linked to task-induced deactivation, our analytic approach departed from prior work: we first identified areas where task-related activity correlated with IID in ANS measures, and then determined whether these areas were associated with task-related activation or deactivation.

We employed a silent mental arithmetic (MA) subtraction task, in absence of any exogenous behavioral performance. The task was conducted in a blocked manner to allow modeling task-induced changes in activity. Then, as a first step, we used a group-level voxel-wise robust-regression approach to identify brain areas where IID in ANS indices correlated with task-induced signal change. These regions then constituted functional regions of interest (fROIS). In a last step, for each fROI, we conducted group-level tests to determine whether it showed statistically significant task-induced activation or deactivation. In this way, our approach did not limit the Brain/ANS investigation to regions that were necessarily strongly (de)activated by the task, while still offering the possibility to determine which ANS-related clusters were activated or deactivated. We used the MA subtraction task because it produces systematic fronto-parietal activations as well as significant deactivations (Grabner, Ansari, Koschutnig, Reishofer, & Ebner, 2013). MA tasks also perturb both heart rate and respiratory rate (Widjaja et al., 2015). Furthermore, IID in ANS responses during MA tasks (heart rate or HRV) correlate with magnitude of EEG or NIRS indicators (Tanida, Sakatani, Takano, & Tagai, 2004; Yu, Zhang, Xie, Wang, & Zhang, 2009).

We examined several independent ANS measurements, ranging from standard indices of heart rate and respiratory variability (Task Force, 1996) to a recently developed information-theoretic measure conditional self-entropy (Widjaja et al., 2015) that quantifies the amount of information (variance) in the cardiac time series after accounting for the effects of respiration, thus allowing more precise characterization of the sympathetic contribution to the cardiac activity.

## 2. Methods

### 2.1 Participants

Thirteen participants (9 males, *Age* = 24.6 ± 3.5) participated in the study. Physiological data of two participants were excessively noisy and so these were not included in the analysis.

### 2.2 Brain imaging acquisitions

We acquired a single structural scan per participant at the beginning of the experimental session, using a 3D T1-weighted Magnetization Prepared RApid Gradient Echo (MPRAGE) sequence (TR/TE = 2700/4.18 ms, flip angle = 7°, voxel size = 1 mm isotropic, matrix = 256×224, 176 sagittal slices). We acquired several functional imaging scans. The first consisted of 115 volumes (TR/TE = 2200 / 33 ms, flip angle = 75°, voxel size = 3×3×3 mm, 0.45 mm slice spacing, matrix = 64 × 64, 37 axial slices covering the entire brain). During this scan an on/off mathematical task was carried out by participants, and this scan is the focus of the current report. The other scans consisted of long resting state scans or long scans where participants continuously engaged in mathematical computations in absence of rest periods. These are not discussed here.

### 2.3 Task and procedure

During the fMRI session, participants engaged in a mental arithmetic Continuous Performance Task (CPT), which had a 4-cycle on/off structure, where the durations of task and rest cycles was 28 and 16 seconds respectively. During the rest periods participants were asked to observe a fixation cross. The run began with 45sec of rest and then continued into the 4 on/off cycles. Each cycle began with a written ‘start’ prompt accompanied by an arithmetic expression, and ended with ‘stop’ prompt. The start cue consisted of an arithmetic expression such as “510 11 = 499” indicating to begin subtracting “11” from the starting number “510”. After two seconds the cue disappeared and participants were instructed to continue subtracting covertly until a STOP instruction appeared on the screen. To avoid excessive practice effects there were 4 different versions of the subtraction task: continuous subtraction of 11, 7, or 13, and another block where participants subtracted 2 and 3 repetitively (e.g., 510 -2 -3 -2 -3…). The task was self paced. The last ‘on’ cycle was followed by an additional 45sec of rest. Thus, the overall duration of the rest and task periods considered consisted of 4 × 28 = 112 seconds and 45+3 × 16+45 = 138 seconds, respectively.

### 2.4 Recording and processing of physiological data

We acquired physiological data during all functional scans. Cardiac and respiration data were acquired using the scanner‘s built-in equipment at an acquisition rate of 50Hz and stored for offline analysis. Cardiac sequences were recorded via a photoplethysmograph (PPG) device placed on participants’ left forefinger. Respiration data were collected using the displacement of a sensor placed on a belt around participants’ chests.

To derive autonomic indices, we started by extracting cardiac beat-to-beat intervals (BBI) time-series from the PPG data (see Fig. 1a,b). BBIs were initially identified using an unsupervised procedure and these were inspected by visual superimposition of the cardiac events onto the original cardiac time-series. In order to detect the correct sequences of heartbeat events, artifacts, missing and ectopic beats were manually annotated and time-series modified accordingly.

**Figure 1:**
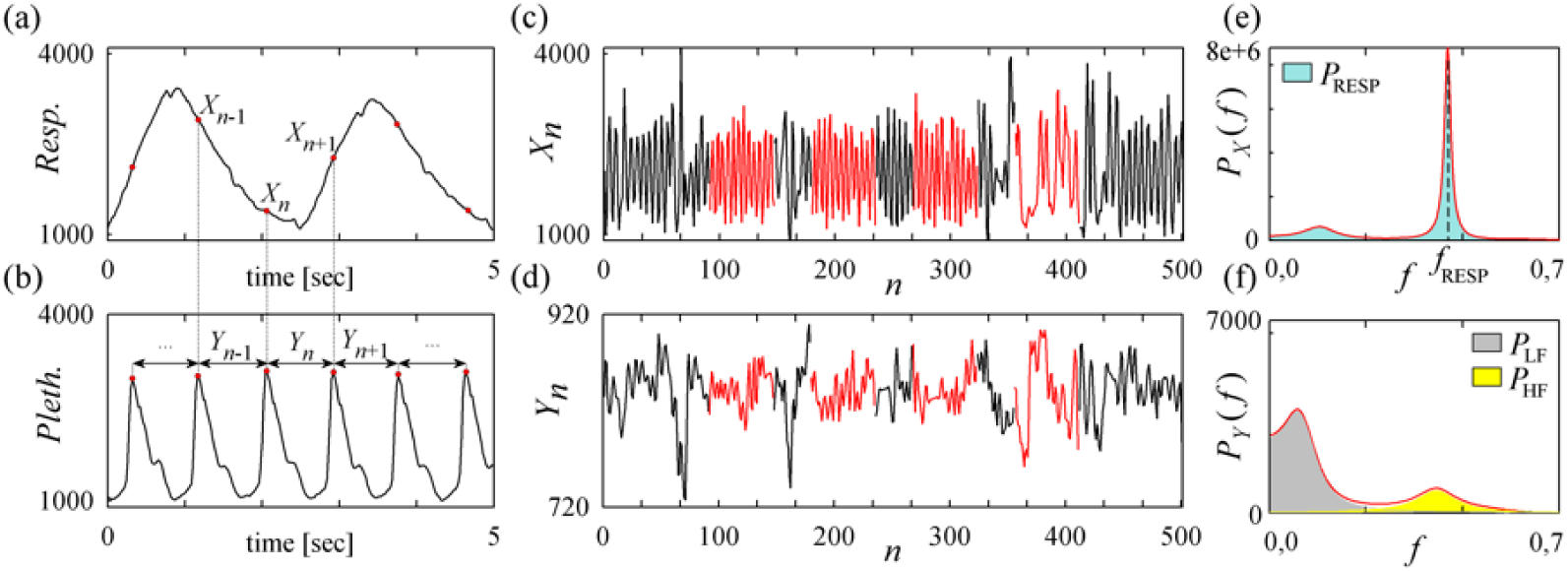
Computation of autonomic indices through univariate parametric spectral analysis. (a) Respiration signal; (b) photoplethysmographic signal and measurement of consecutive cardiac beat-to-beat intervals; (c) respiration series resampled to 2 Hz; (d) series of the cardiac BBIs [higher values indicate a longer break between two successive beats] resampled to 2 Hz; (e,f) power spectral densities of the portions of the respiration and cardiac time series measured during task (i.e., these are the portions of the time series depicted in red in (c,d)), evidencing respiratory frequency and power, as well as low and high frequency power of BBI. Note that the estimation of autoregressive parameters allows to characterize in terms of frequency and power the spectral content of the cardiac series, with strong power in low frequencies (gray), and a spectral peak centered exactly at the respiration frequency (yellow).

The first two indices we derived were mean BBI and RMSSD, standard indices commonly used to assess the autonomic function from heart rate data:

1. The mean *Beat-to-Beat-Interval* (BBI): this is simply the average of the BBIs during the task period. Lower BBI values mean faster heart rate.
2. Root Mean Square of Subsequent Differences (RMSSD) of the BBIs: a time-domain measure (see Berntson, Lozano, & Chen, 2005, for review) that reflects a mainly vagal HRV component, and is thought to capture respiratory sinus arrhythmia. It correlates with power in higher frequencies of the heart period. It was computed as in Eq. 1 where *N* is the number of measured BBIs. RMSSD measures over short time periods (as low as 10sec) have been shown to be good surrogates of much longer recordings (Wang & Huang, 2012). ^1^

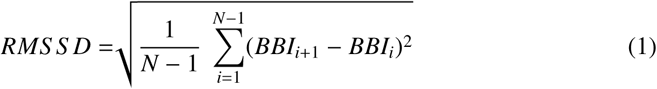

The BBI and respiration time series were interpolated and resampled uniformly at 2 Hz (Task Force Of The European Society of Cardiology, 1996) using cubic spline interpolation. Prior to further analysis, the two time series were de-trended using a zero-phase high-pass filter (IIR, order 2, cutoff frequency=0.0215 Hz) in order to foster the fulfillment of stationarity criteria. Stationarity of each time series was carefully checked through visual inspection. Then, the two time series were analyzed in the framework of autoregressive (AR) modeling, a well-known method for the time- and frequency-domain description of cardiorespiratory time series (Task Force, 1996), widely employed also in cognitive studies (Lane et al., 2009; G. Park, Van Bavel, Vasey, & Thayer, 2013; Williams et al., 2015). Specifically, the time series of respiration (series or BBI (series *Y*) were first described individually using the univariate AR models in Eq 2 where *X*_*n*_ and *Y*_*n*_ are the *n*-th samples of the time series, *p* is the model order, *A*_*k*_, *B*_*k*_ are linear regression coefficients defined for each *k* = 1, …, *p*, and *U*, *V* are the time series of the model residuals. Note that the order *p* was estimated separately for each participant using a BIC criterion (*Mean* = 6.4 ± 0.8; see Schwarz, 1978).

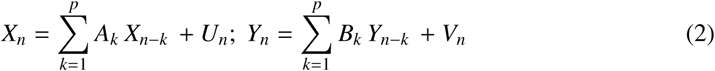

We used a well-established procedure for deriving the frequency content of the cardiac and respiration time series from the estimated AR parameters (Baselli, Porta, Rimoldi, Pagani, & Cerutti, 1997). Exploiting the frequency-domain representation of the models in Eq. 2 we computed the power spectral density of each series, denoted as *P*_*X*_(*f*) and *P*_*Y*_ (*f*), quantifying the power of *X* and *Y* as a function of frequency (Baselli et al., 1997). As also seen in the example in Fig. 1, this representation was used to compute indices related to the total power of the time series, the power confined to specific frequency bands, or the frequency of specific oscillatory components, as we detail below.

To identify the part of heart rate variability that was unrelated to respiration we performed joint analysis of the cardiac and respiration time series using the two bivariate models in Eq. 3, respectively describing the dependence of the current cardiac BBI on the past respiration values (upper model) and its dependence on both its own past values and the past respiration values (lower model). The differential predictive ability of the two models in Eq. 3 is expressed by the variance of the model residuals *W* and *Z*, denoted respectively as Σ_*W*_ and Σ_*Z*_. In this formulation, an index of the ability of the past BBIs to predict the present interval above and beyond the predictability provided by the past respiration samples is seen in the conditional self-entropy (cSE), an information-theoretic measure defined as *S*_*Y*|*X*_ = 0.5 *ln*(Σ_*W*_ /Σ_*Z*_) (Faes, Porta, & Nollo, 2015). The cSE captures the predictability of the time series *Y* based on its own past, and is unaffected by the strength of the influences exerted on *Y* by the other modeled series *X*.

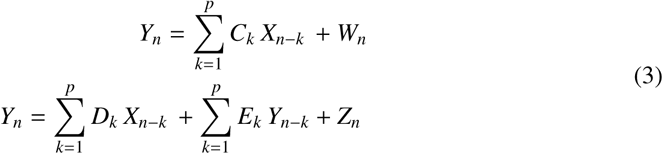

The methodology described above was applied, separately, to the portions of the BBI and respiration time series corresponding to the ‘on’ and ‘off’ periods of each participant‘s task (by censoring time points from the regression model). Specifically, by assuming stationarity across periods belonging to the same condition (i.e., task vs. rest), we obtained realizations of Eqs 2 and 3 drawing on present and past points of the two time series separately from the rest periods or the task periods. We finally considered, respectively, the 276 and 224 time series samples for each participant derived as described in Section 2.3. From these realizations, univariate and bivariate AR models were identified using the standard least squares approach, and estimating the model order according to the Bayesian Information Criterion (Faes, Erla, & Nollo, 2012). The estimated model coefficients were used to derive, for each participant, two values (one for the ‘off’ periods and one for the ‘on’ periods) for three frequency related autonomic indices (see Fig. 1 for an example): Peak Frequency and Power of Respiration and Low-to-high frequency power ratio of heart rate variability.

In all, using these procedures we derived the following six autonomic indices: the first three extracted from time-domain analyses and the last three from frequency domain analyses.

1. (Time Domain) *Mean beat-to-beat interval* (BBI): The average of the BBIs during the task-ON period
2. (Time Domain) *Root Mean Square of Subsequent Differences* (RMSSD) of the BBIs: to identify high-frequency signatures of the cardiac periods, reflecting a mainly vagal HRV component;
3. (Time Domain) *Conditional Self Entropy* (cSE): this measure is computed as explained above from the error variances of the bivariate AR representation of BBI and respiratory time series;
4. (Frequency Domain) *Power of respiration* (*P*_*RES* *P*_): the measure is obtained as the total power of the respiratory time series computed as the area under the spectral profile *P*_*X*_(*f*);
5. (Frequency Domain) peak *Frequency of respiration* (*f*_*RES P*_): the measure corresponds to the frequency of the main oscillatory component of respiration, assessed for the main spectral peak;
6. (Frequency Domain) *Low-to-high frequency ratio of heart rate variability* (*P*_*LF*_ /*P*_*HF*_): this measure is computed as the ratio between the power of low (LF, 0.04-0.15 Hz) and high frequency (HF, 0.15-0.4 Hz) components of the BBIs, where each power is assessed as the area under the profile of the spectral components located in the LF or HF band.

The first two indices constitute a basic approach to autonomic acquisitions. Moreover, given that heart rate variability also reflects respiration patterns, we introduced cSE to characterize the extent to which the BBI series is uniquely predicted by its own past, as described above(see also, Widjaja et al., 2015). We derived indices 4 and 5 because mental stress impacts the amplitude (spectral power) and frequency of respiration (Masaoka & Homma, 1997). Index 6, the LF/HF ratio computed for the BBI series, is a common measure for assessing the sympathovagal balance (Task Force Of The European Society of Cardiology, 1996), which is also known to be affected by stress (Pagani et al., 1991). In addition, to maintain consistency with prior literature we calculated the non-normalized power in high and low frequencies of the cardiac series. The correlations of these with RMSSD and LF/HF ratio were 0.27 and 0.15 for PHF and 0.06 and 0.71 for PLF. Results for these measures are shown in supplementary materials.

### 2.5 Neuroimaging analysis

#### 2.5.1 Pre-processing

We implemented the following neuroimaging pipeline. We discarded the first 14 acquired volumes (30.8 seconds) to allow for steady-state magnetization delay. Pre-processing was performed using AFNI (Cox, 1996). Time series were de-spiked, corrected for slice-timing differences and spatially aligned to a reference acquisition. We applied spatial smoothing using a tridimensional Gaussian with fwhm of 6mm^3^

#### 2.5.2 Deriving task-related activation at the single participant level

On the single-subject level, time series analysis was performed using multiple-regression methods implemented via AFNI‘s 3dDeconvolve utility. There were three regressors related to the experimental design: one capturing the on/off task structure, and two capturing the Start/Stop prompts at the beginning and end of each block. These timings were convolved with a boxcar hemodynamic response function. Additional regressors were the 6 motion parameters estimated from the alignment procedure. The Beta estimate of the regressor for the on/off task structure was the one of theoretical interest and propagated to a group-level analysis as described below.

We projected the single-participant statistical maps into 2D-surfaces using a Freesurfer (Fischl, Sereno, Tootell, & Dale, 1999) and SUMA (Saad & Reynolds, 2012) processing pipeline as follows: the single participants’ structural images were aligned to the first image of the first functional run (to maximize T1 effect in the EPI data) and alignment was manually checked and adjusted when needed. The next steps were implemented using FreeSurfer‘s procedures. Here, the anatomical volumes were processed using a pipeline in which they were registered to a reference space, segmented, skull stripped, cortex-extracted and inflated to a 2-dimensional cortical-fold representation. This cortical representation was registered to common space using FreeSurfer‘s registration procedures. It was in this 2D cortical domain where all statistical analyses were conducted.

#### 2.5.3 Group-level analysis

Before proceeding to our main analysis of ANS correlates, we first examined whether, at the group level, the task produced activity patterns similar to those found in prior studies. We projected activation maps from the single-participant level to each participants cortical surface representation, and all group level analyses were performed on the cortical surface. We constructed typical activation maps by testing participants Beta values at each vertex against 0. In this analysis, statistical significance was set at *p* < .005 (uncorrected single-voxel cluster forming threshold), and corrected for family-wise error (*p* < .05) via cluster-wise thresholding, which identifies contiguous clusters of statistically significant voxels (FWE *p* < .05 using cluster extent).

We implemented cluster-based correction on the 2D cortical surface rather than the 3D volume, as all analyses were conducted on the surface. The general framework was as follows. We generated random data from a normal distribution, smoothed those with the input-based spatial-autocorrelation estimate as detailed below, identified the top 1% of voxels (our uncorrected single vertex level), clustered those and stored the value of the largest cluster. Storing the top 1% was done to match our analysis procedure, which identified clusters where all correlation values had to have the same sign (i.e., we split the correlation map into positive and negative values prior to clustering). We derived the smoothness estimates for the Monte-Carlo simulation from the single-participant spatial auto-correlation of the task-based residuals, after projecting those to the surface. This procedure might *over*-estimate the ‘null’ spatial autocorrelation, as the task-analysis regression that produces those likely fails to remove all task-related activation. This will maintain structure in the residuals, thus increasing spatial autocorrelation. This produced an estimate of 9mm smoothness on the surface.

Our main analysis was focused on identifying areas where the Beta values correlated with inter-individual differences in autonomic measures. To this end we conducted a robust regression analyses on the single vertex level (using the *lmrob* function of the *robustbase* package in R) where the set of Beta values for the vertex was the predicted variable and the autonomic measure the predicting variable. This regression returned a significance value for each regressor. Family-wise error correction was implemented as described above (single voxel *p* < .01 uncorrected; FWE controlled using cluster extent, *p* < .05). We could thus identify clusters where *all vertices* showed a significant correlation between the Beta values and the autonomic measure (a significant brain/behavior correlation).

In a last step, we treated each cluster in which the brain/behavior correlation was significant as a functional region of interest (fROI). For each fROI we determined if it was associated with task-related activation or deactivation, by calculating the mean β in the cluster per participant, and then submitting these values to a group-level T-test against 0. These latter tests were FDR-corrected within each autonomic measure, effectively controlling for the number of clusters showing significant Beta/ANS correlations.

## 3. Results

### 3.1 Autonomic indices during task periods and between-task intervals

The values for the different ANS indices during task performance were as follows: BBI:*M* = 0.86*s* ± 11, range: 0.68–1.06; RMSSD: *M* = 47*ms* ± 17, range: 26–71; pRESP:*M* = 381805 ± 83514, range: 271451–548121; fRESP: *M* = 0.32*Hz* ± 0.07, range: 0.18–0.43;LF/HF: *M* = 2.19 ± 1.96, range: 0.3–7.08; cSE: *M* = 1.01 ± 0.31, range: 0.49–1.49.

Prior to analyzing the autonomic indices we evaluated all pair-wise correlations between the 6 measures during task performance. As shown in Figure 2, correlations were relatively moderate, with the maximal correlation holding between cSE and BBI, Pearson‘s *R* = 0.55. For this reason we correlated each measure separately against BOLD activity, rather than using partial correlations. We also evaluated the correlation of these measures with age of our participants (in months). For BBI, RMSSD and cSE, the absolute correlation value was lower than 0.1. For pRESP it was 0.16, for fRESP it was -0.35 and for LF/HF it was 0.52. None approached significance.

**Figure 2:**
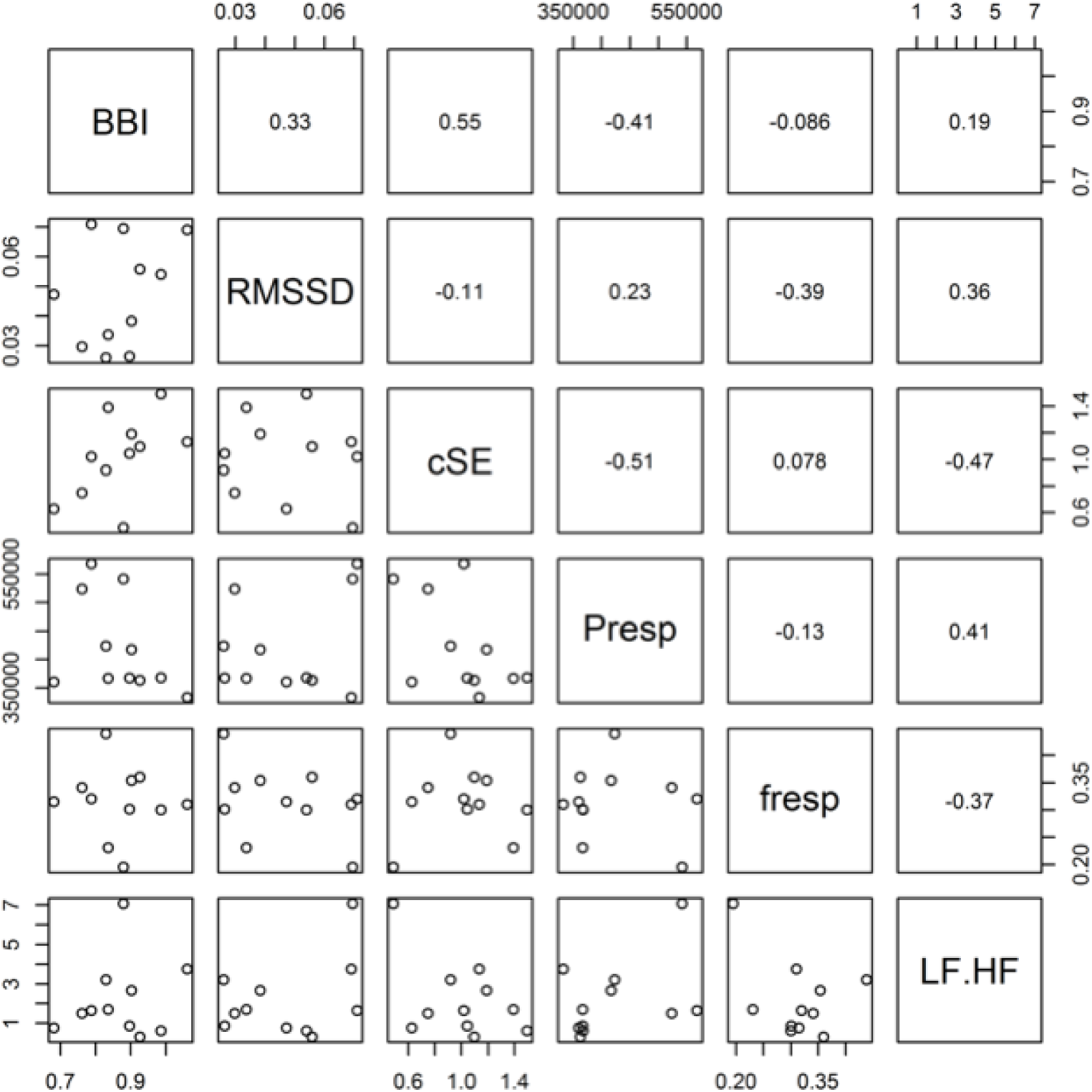
Strength of correlations between autonomic measures computed for each individual participant during task execution.

To evaluate task effects we examined differences in autonomic activity during the task-on and task-off periods. We replicated prior results (e.g., Taelman, Vandeput, Vlemincx, Spaepen, & Van Huffel, 2011) showing reduced RMSSD values during the math task as compared to rest (twotailed T-tests); RMSSD: *t*(10) =–2.20, *p* = .05. This points to lower vagal modulation during task performance. The LF/HF ratio also differed significantly, in the same direction reported in prior work; *t*(10) =–2.69, *p* = .02. The difference in BBI was not significant. For fRESP, pRESP and cSE the results were marginal: For fRESP: *t*(10) = 1.60, *p* = .07, with slightly higher values for on than off period (*M* = 0.32± 0.02 vs. 0.28 ±0.02); for pRESP values were conversely greater for the off period, *t*(10) = 1.49, *p* = .083; cSE was marginally higher during the off than on period, CSE: *t*(10) = 1.69, *p* = .06.

We could not directly address the reproducibility of the different ANS measures via common test-retest procedures, because the ANS indices are likely to vary across blocks due to practice or attention-related effects. However, we could perform another type of analysis. Specifically, because we derived these measures for the rest periods between blocks we could quantify to what extent inter-individual differences in the ANS measures maintained across task and rest epochs. The resulting (Pearsons) correlations were as follows: BBI (*R* = 0.98, *p* < .001); RMMSD (*R* = 0.87, *p* < .001); LF/HF (*R* = 0.96, *p* < .001); pRESP (*R* = 0.57, *p* = .067); fRESP (*R* = 0.66, *p* = .027); cSE (*R* = 0.92, *p* < .001).

### 3.2 Task-induced activation and deactivation

Consistent with prior studies of mental arithmetic, we found task-related activation in areas involved in attention and verbal rehearsal (e.g., left inferior frontal gyrus), and fronto-parietal regions. The distribution of deactive regions formed a good match the topology of the Default Mode Network (Figure 3).

**Figure 3:**
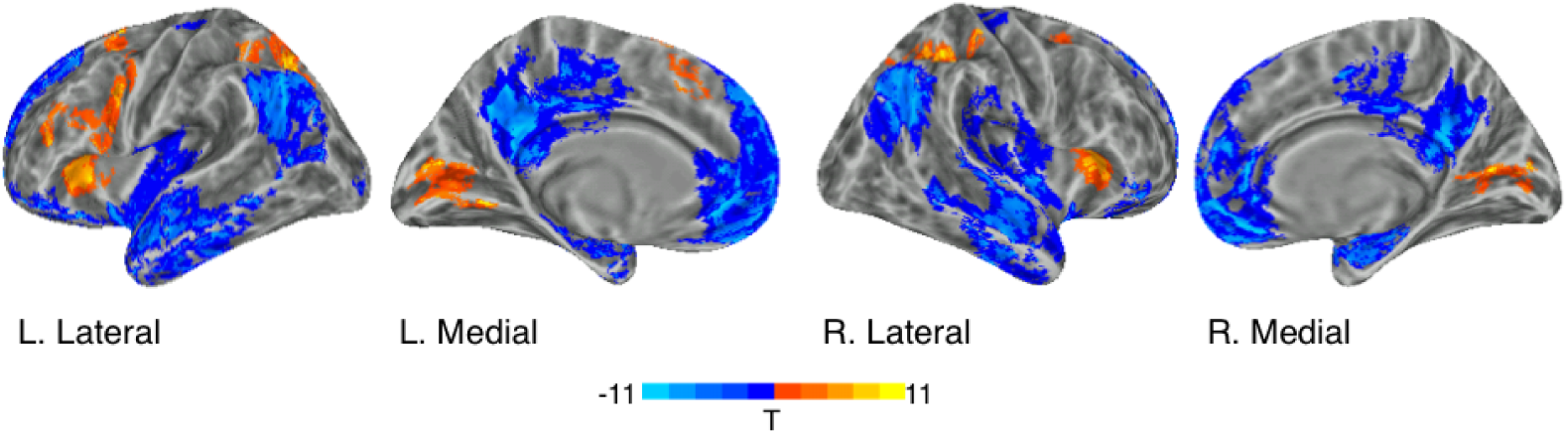
Task-induced activation and deactivation during the 4-block mental arithmetic task. The task consisted of four task-on (28s)/task-off (16s) cycles.

### 3.3 Correlations between task-invoked activity and autonomic indices

#### 3.3.1 Beat-to-Beat Interval (BBI)

Prefacing the specific findings, we found that individuals with lower heart rate (higher BBI) showed more moderate task-induced activation in fronto-parietal regions, as well as more moderate task-induced deactivation in vmPFC.

Specifically, we found positive BOLD/BBI correlations in the left occipitotemporal cortex (Figure 4A). On the right (Figure 4C), we identified three clusters consisting of the STS (#1) and ventro-medial prefrontal cortex (#2, 3). For the more inferior vmPFC cluster (#3) we found significant deactivation on the group level (FDR corrected for 6 tests). Taken together with the positive correlation, this means that individuals with a higher-value BBI (slower heart rate) showed less deactivation in vmPFC.

**Figure 4:**
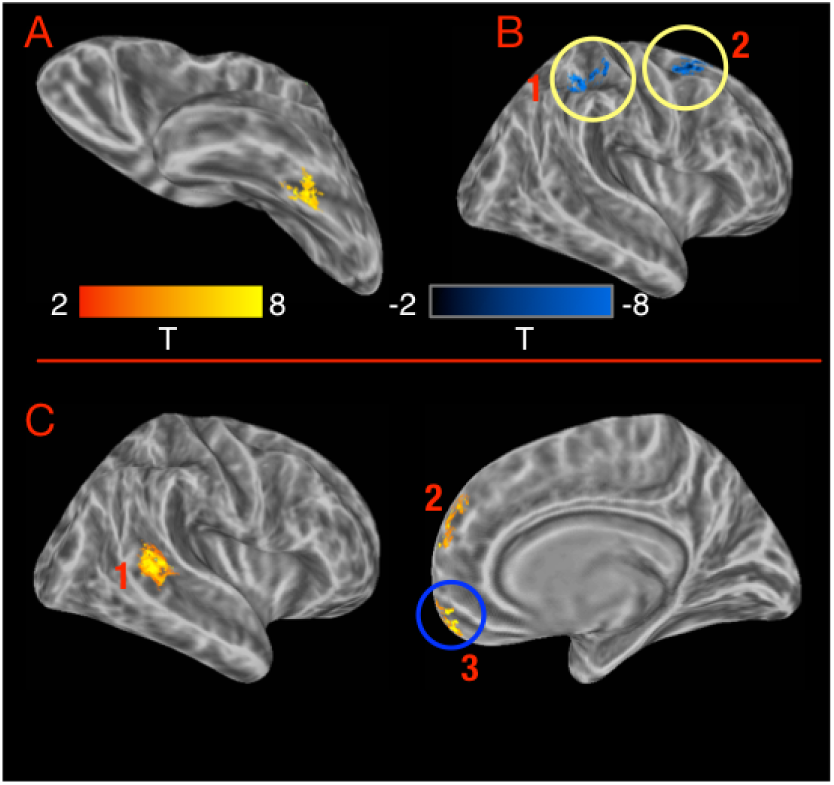
Interindividual differences in BBI. Panel A: Left hemisphere region showing positive BBI/BOLD correlations. Panel B: Right hemisphere regions showing negative BBI/BOLD correlations. Panel C: Right hemisphere regions showing positive BBI/BOLD correlations. Here and in all subsequent figures, Yellow circles mark correlation clusters that showed statistically significant taskrelated activation and Blue circles mark correlation clusters that showed statistical significant task-related deactivation. Note that negative correlation indicates that greater activity was linked to higher heart rate (shorter BBI).

Negative BOLD/BBI correlations were found in two clusters on the right (Figure 4B). These were found around IPS (#1) and SFS (#2). In both clusters mean group-level activity was significantly *above* baseline (FDR corrected for 6 tests). This means that individuals with a higher BBI (slower heart rate) showed weaker above-baseline activation in these regions. Note that these clusters match regions that were identified as particularly strong above-baseline activity in the initial task analysis (Figure 3).

#### 3.3.2 Conditional self-entropy (cSE)

Several brain regions showed a negative BOLD/cSE relationship (Figure 5). These were located, bilaterally, in the superior frontal gyrus (SFG) and the superior anterior insula. A fifth cluster was found in the central cingulate gyrus. Higher cSE reflects better predictability of the BBI based on its recent past after discounting for the respiratory effect, which is generally associated with the rise of more regular HRV oscillations related to higher sympathetic tone.

**Figure 5:**
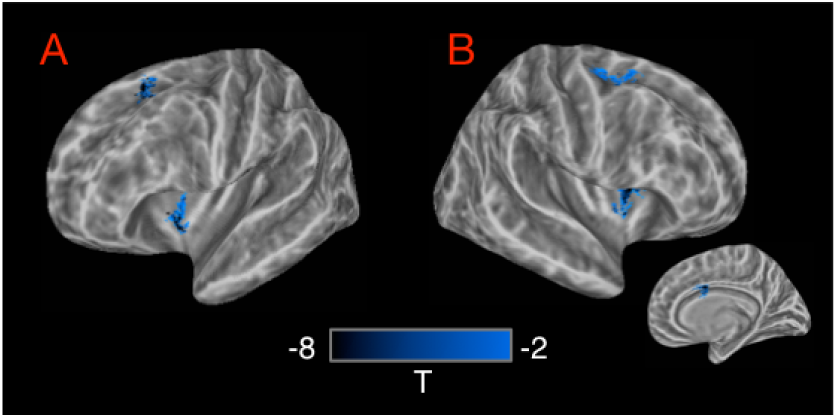
Interindividual differences in CSE. Panel A, B: Left and Right hemispheres. Inset: cingulate gyrus on medial surface of right hemisphere.

#### 3.3.3 RMSSD

We found an extended bilateral network of regions showing positive correlations between task-induced activity and RMSSD (Figure 6). No negative correlations were found (Figure 6A,B). In 18 of the 29 identified clusters, task-related changes were associated with significant deactivation (FDR corrected for 29 tests; see Figure 6C), and none showed significant activation. In summary, increased RMSSD in these regions, which overlapped substantially with the Default Mode Network, was associated with less marked deactivation patterns.

**Figure 6:**
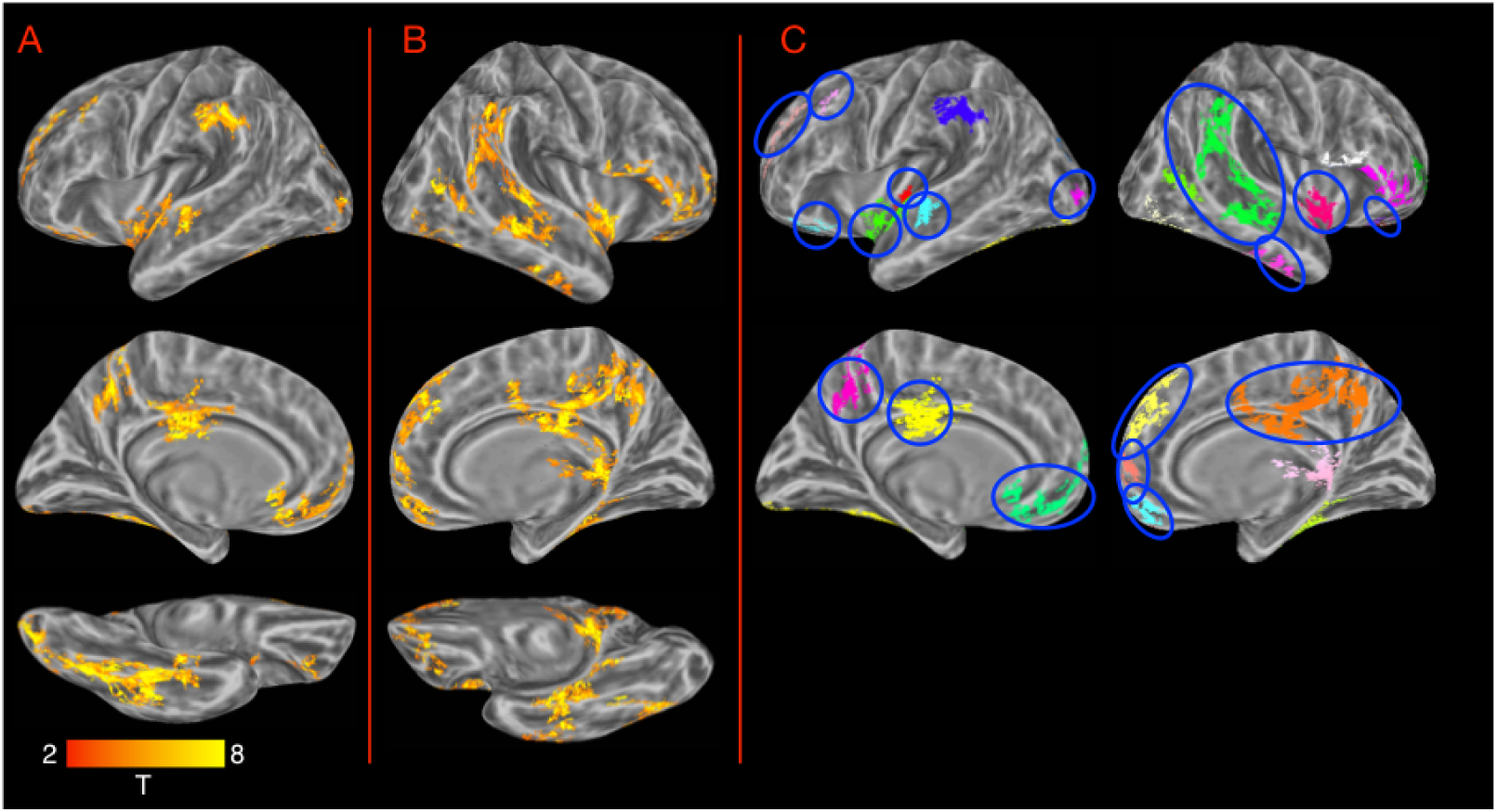
Interindividual differences in RMSSD. Panels A, B: Left and Right hemisphere regions where task-related activity correlated with RMSSD. In all cases the correlations were positive. Panel C: depiction of separate clusters marking those clusters that showed significant deactivation (FDR corrected for 29 tests) at the group level. No cluster showed significant activation.

In the left hemisphere (Figure 6A) these regions included (in 13 clusters) the SMG, anterior STG extending into the supratemoral plane (STP), SFG, anterior left IFG, midline regions including the precuneus, the central part of cingulate gyrus, rostral part of ACC and vmPFC. There was also an extensive cluster in the occipitotemporal cortex. On the right (Figure 6B), these areas included (in 16 clusters) the STS extending posteriorly to the SMG, but also to the MTG, the anterior insula and the right IFG. Correlations were also found in the most anterior part of vmPFC, the precuneus, the central part of the cingulate gyrus, and parahippocampal gyrus.

#### 3.3.4 Low-frequency to High-frequency ratio

Positive BOLD/LF-HF correlations were found in the insula bilaterally, along the posterior, middle and anterior cingulate gyrus, and in a few additional fronto-parietal clusters (Figure 7). No negative clusters were found. In left occipto-temporal cortex, left STS and left orbitofrontal cortex there was statistically-significant deactivation, indicating that participants with greater LF/HF ratio showed weaker deactivation. No cluster showed statistically significant activation.

**Figure 7:**
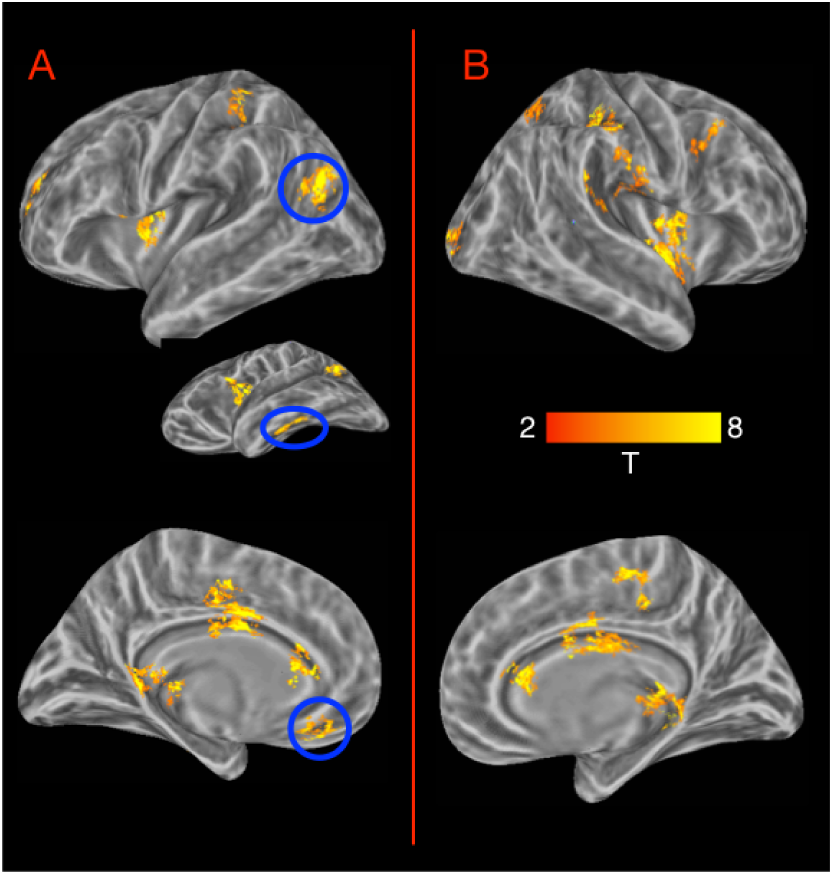
Interindividual differences LF/HF ratio. Panels A, B: Left and Right hemisphere regions where task-related activity correlated with LF/HF ratio. In all cases the correlations were positive. Blue circles mark clusters showing significant deactivation at the group level (FDR corrected for 18 tests).

#### 3.3.5 Respiration frequency (fRESP)

Respiration frequency (fRESP) was associated with task-induced responses both positively and negatively (22 clusters in all; Figure 8). Generally, increased fRESP was linked to greater activation in task-active regions and greater deactivation in task-deactive regions.

**Figure 8:**
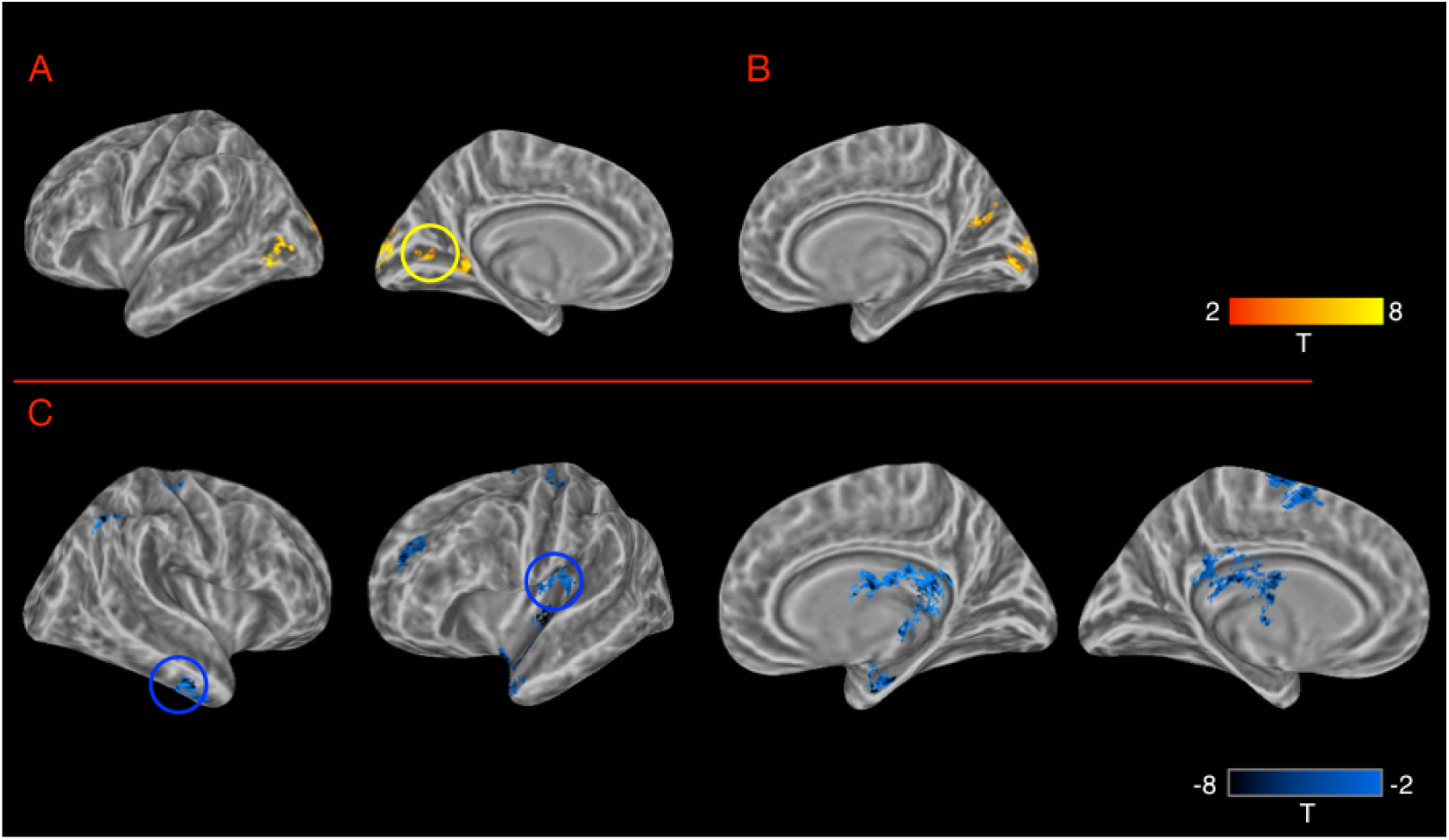
Interindividual differences in respiration frequency (fRESP). Panels A, B: Left and Right hemisphere regions where task-related activity correlated positively with fRESP. One region (yellow circle) showed significant taskrelated activation (FDR corrected for 27 tests). Panel C: regions where task-related activity correlated negatively with fRESP. Blue circles mark clusters with statistically significant task-related deactivation (FDR corrected).

Positive BOLD/fRESP correlations were found, on the right (Figure 8B), in the calcarine sulcus and nearby cuneus, and the posterior parietal-occipital fissure, and within the calcarine sulcus on the left (Figure 8A). This latter cluster showed significant above-baseline activation meaning that greater fRESP was associated with greater activation. Indeed, as can be seen in the task-activation map (Figure 3), the occipital regions identified here formed a good match with the posterior midline clusters showing significant task-activation in the whole-brain analysis.

Negative BOLD/fRESP correlations (Figure 8C) were found, bilaterally, in central insula, and posterior cingulate regions. On the left, other clusters were found in SFS. On the right, clusters were found in posterior PoCG and anterior PHG. Several of these clusters showed significant task-induced deactivation (FDR corrected for 27 tests). There was no indication for above-baseline activation in any of the clusters showing negative correlations.

#### 3.3.6 Respiration power (pRESP)

Respiration power (pRESP) was associated with task-evoked responses mainly in sensorimotor and midline regions. These clusters largely excluded the lateral frontal cortex, and very few were found in parietal and occipital cortices. With one exception, correlations were positive, but as shown in Figure 9, in many of these clusters, the task produced significant deactivation (activations were not found). Thus, generally, greater respiration power was associated with less deactivation.

**Figure 9:**
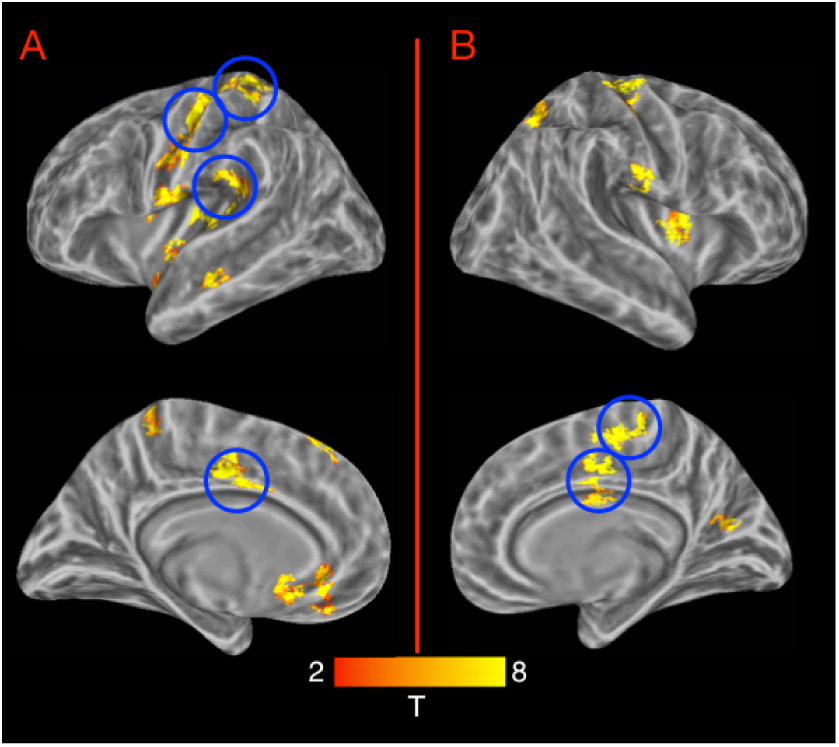
Interindividual differences in respiration power (pRESP). Panels A, B: Left and Right hemisphere regions where task-related activity correlated with pRESP. In all cases the correlations were positive. Blue circles mark clusters with statistically significant task-related deactivation (FDR corrected).

In both hemispheres we found correlations in the superior temporal plane and insula (posterior and central). Additional correlations were found in left STS and the right central sulcus. Statistically significant deactivations (FDR corrected for 5 tests) were found in midline regions, and left parietal operculum. A single cluster in the right posterior insula (not presented in Figure 9) showed a negative correlation with task-induced activity.

### 3.4. Summary of results

The above-presented findings can be summarized by considering the relationship between areas showing BOLD/ANS relations and those that show task-related activation or deactivation at the group level (i.e., regions shown in Figure 3). It is also possible to quantify, for each autonomic index, the distribution of areas that were significantly task-activated or deactivated, as evaluated separately within the functional ROIs showing BOLD/ANS correlations.

Figure 10 shows the overlap between task-related effects and BOLD/ANS effects. As can be seen, BOLD/ANS correlations were found in core nodes of the DMN, but sparing the PCC. There was less overlap with task-active regions, most notably for those in left dorsolateral prefrontal cortex.

**Figure 10:**
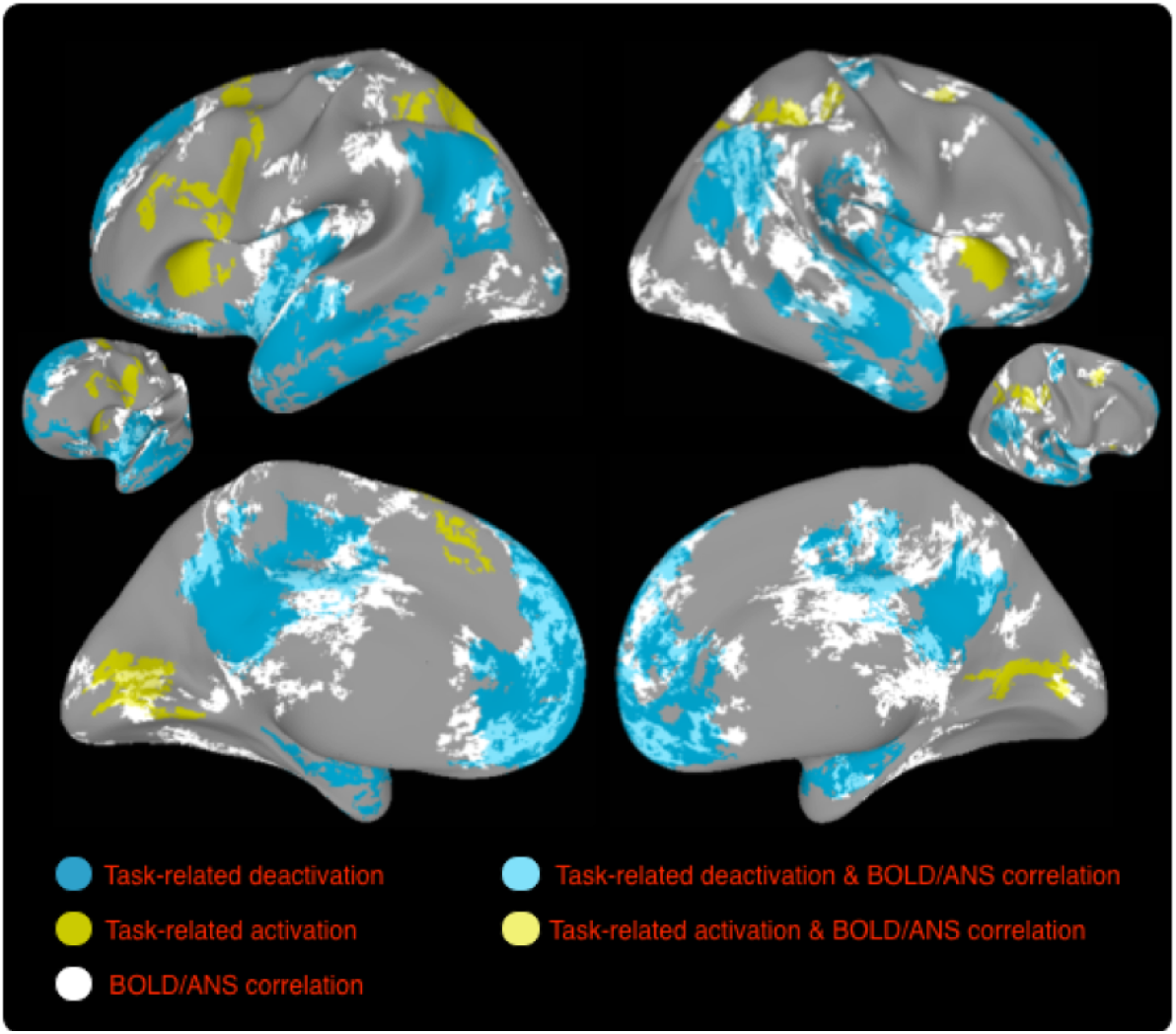
The figure shows in yellow and blue areas of activation and deactivation during the mental arithmetic task. Areas showing BOLD/ANS correlations for any of the autonomic measures, are shown in white.

Table 1 shows, for each autonomic measure, the proportion of brain surface area linked to statistically significant task-related activation, task-related deactivation, or no difference from baseline (proportions sum to 100% within row). This analysis shows that by and large, interindividual differences in ANS activity were more related to task-related deactivation than to activation. (These proportions hold for the particular single-voxel cluster-forming threshold we used, and different results may be obtained for other cluster-forming thresholds.)

**Table 1:** Task effects in areas identified by BOLD/ANS analyses. For each measure, the relative area of significantly active, significantly deactive and other clusters is shown (each row sums to 100%)

| Measure | Positive BOLD/ANS correlation |  |  | Negative BOLD/ANS correlation |  |  |
| --- | --- | --- | --- | --- | --- | --- |
|  | % Act. | % Deact. | % Null | % Act. | % Deact. | % Null |
| BBI | 0 | 9 | 46 | 45 | 0 | 0 |
| cSE | 0 | 0 | 0 | 0 | 0 | 100 |
| RMSSD | 0 | 63 | 37 | 0 | 0 | 0 |
| fRESP | 4 | 0 | 35 | 0 | 26 | 35 |
| LF:HF | 0 | 11 | 87 | 0 | 0 | 2 |
| pRESP | 0 | 72 | 25 | 3 | 0 | 0 |

## 4. Discussion

In the current study, participants performed a continuous mental arithmetic task known to induce mental effort (Tanida et al., 2004; Widjaja et al., 2015; Yu et al., 2009). We replicated task-related activation and deactivation patterns previously reported for this task (e.g., Grabner, Ansari, Koschutnig, Reishofer, & Ebner, 2013). We also replicated prior findings of reduced cardiac RMSSD during task execution (e.g., Duschek, Muckenthaler, Werner, & del Paso, 2009).

In tandem, we found that inter-individual differences (IID) in ANS measures were associated with task-induced deactivation. In particular, the strength of task-induced deactivation was significantly negatively correlated with indicators of the vagal parasympathetic ANS control, such as the mean and RMSSD of the cardiac BBIs and the respiratory power and frequency. Equally important, different ANS measures were associated with deactivation in different regions, suggesting they load on different latent constructs underlying deactivation. Finally, ANS measures were less-extensively linked to task-induced activation.

In what follows we address potential accounts for these correlations, particularly for the relation between IID in ANS measures and deactivation. We do so in relation to what is currently known about the computations that these regions mediate during mental arithmetic. We then discuss the implications of these findings for research documenting a “failure to deactivate” in certain populations, and for studies interested in quantifying task-related activations (or deactivations) more generally.

### 4.1 Activation during mental arithmetic

As a result of several neuroimaging studies of mental subtraction, there exists a good basis for interpreting functions associated regions that are activated or deactivated during this task. As opposed to numerical multiplication, which involves extensive access to episodic knowledge, subtraction appears to rely on numerosity comparison mechanisms (Prado et al., 2011), and may load on both phonological working memory and spatial representations (Cavdaroglu & Knops, 2016; Kallai, Schunn, & Fiez, 2012). A set of fronto-parietal and few occipitotemporal regions have been consistently implicated in this task. For instance, parietal regions (right superior), prefrontal regions and the fusiform bilaterally have been linked to abstract-level arithmetic computations in a paradigm where arithmetic expressions were followed by visual dot patterns whose numerosity matched or did not match the correct sum of the expression (Kallai et al., 2012), suggesting these regions code for abstract numerical quantities. Others have documented similar fronto-parietal activations (Grabner et al., 2013; Vansteensel et al., 2014). For instance, using ECoG and fMRI, Vansteensel et al. (2014) implicated an area straddling the junction of the precentral gyrus and middle-frontal gyrus in the implementation of subtraction. They presented one problem at a time and found that this region showed a continuous response throughout the trial including the period following the removal of the arithmetic problem from the screen. The fusiform gyrus similarly showed a strong initial response (corresponding to processing the visual stimulus), which continued to be elevated during the computation time. Thus, frontal and occipital task-related activation appears to be related to the subtraction computation itself.

### 4.2 Deactivation during mental arithmetic

Particularly relevant to our investigation are brain areas that show deactivation during mental arithmetic. Prior work suggests that during mental arithmetic, areas showing deactivation are implicated in the arithmetic computations themselves. Grabner et al. reported deactivation patterns extremely similar to the ones we found. Notably, they found that the magnitude of deactivation in some of these areas (angular gyrus [AG] bilaterally and ACC) was stronger for larger problems. In addition, both the left AG and anterior left SFG/MFG showed overall deactivation, while accompanied by relatively greater activity for more confusing problems. The latter finding suggests that during mental arithmetic, deactive regions are less deactive for more complex contexts, and most generally indicate that deactive-regions may be involved in relevant computations. This possibility is also supported by EEG and EEG/fMRI studies. In an EEG/fMRI study using source-localization (Sammer et al., 2007), theta activity during mental arithmetic correlated with BOLD responses in many brain regions showing deactivation in our study, including the superior temporal plane and nearby perisylvian regions bilaterally, and both posterior and anterior cingulate gyrus (see also Grabner et al.‘s Figure2B, p. 797). This suggests that deactivation during mental arithmetic cannot be explained in terms of shutting down of a “default” process, but that activity in these regions tracks rapid fluctuations in working memory demands during the task (see also Wager et al. 2009 for link between deactivation and rapid ANS fluctuations). An MEG study (Ishii et al., 2014) also examined sources of theta activity during mental arithmetic and identified frontal midline regions as generators, suggesting that they are linked to the task‘s working memory demands. Meltzer et al. (2007) reported that IID in task-evoked frontal-midline theta during working maintenance correlate with IID in task-evoked BOLD during the same task. In that study, participants with higher theta showed lower BOLD activity in anterior medial prefrontal cortex, inferior parietal lobule and left MFG. As discussed by Meltzer et al., one of the puzzling aspects of the theta/BOLD relationship is that it is negative (i.e., stronger theta accompanied by lower BOLD), even though theta is typically considered to be meaningfully related to neural activity. One possibility they mention is that theta activity is inhibitory in nature, so that increased theta is associated with reduced metabolic demands. Altogether, prior work suggests that (at least some) deactivations during mental arithmetic are linked to increased theta, which in turn is specifically linked to rapidly fluctuating working memory demands of the task.

Less is known on the sources of deactivation of auditory cortex and the superior temporal plane during mental arithmetic. Zarnhofer et al. (2012) found that a multiplication task was associated with deactivation of the left transverse temporal gyrus, a, with greater deactivation associated with less self-reported use of verbalization in that task. There is also some evidence that working memory demands produce deactivation of both auditory and visual sensory cortices (Azulay, Striem, & Amedi, 2009), but we note that in the current study, deactivations were largely limited to auditory cortex.

### 4.3 Task-induced changes and inter-individual differences in ANS metrics

In the current study, task execution was associated with changes to ANS cardiac profiles identified in prior work, most notably producing a reduction in the fast short-term variability of the heart rate, here seen in a significant reduction in the RMSSD values (as shown previously, e.g., Duschek et al., 2009). In addition, we found that task-related activity was significantly correlated with RMSSD in many regions considered part of the DMN. Given that the most extensive pattern of inter-individual effects was seen for RMSSD, and given that it was robustly impacted by task performance (Section 3.1), we first discuss the RMSSD results, and then address the other measures.

#### 4.3.1 Deactivation and RMSSD

RMSSD is an autonomic index that largely loads on the high frequency fluctuation of the cardiac BBI, and is related to vagal parasympathetic contribution; i.e., relatively short-term, rapidly fluctuating effects (though also influenced by sympathetic activity; Berntson et al., 2005). During cognitive tasks, cardiac complexity reduces as a result of a sympathetic activation generally associated with vagal deactivation (Vrijkotte, van Doornen, & de Geus, 2000; Widjaja et al., 2015). This cardiovascular reaction is also documented in our study by a reduction of indices reflecting vagal ANS control (cardiac RMSSD, respiratory power) for participants showing larger task-related BOLD deactivation. In particular, we found that lower RMSSD was associated with lower activation in posterior, central and anterior cingulate gyrus as well as the angular gyrus bilaterally, STS, left SFG, the central insula bilaterally (Figure 6). Almost all these regions showed significant task-related deactivation at the group level. This suggests that maintenance of baseline activity in these regions indicates a weaker cardiovascular response to the task.

Lower cardiovascular involvement may be beneficial for task performance. For instance, in a study (Duschek et al., 2009) of the relationship between task performance and respiratory sinus arrhythmia (RSA; quantified as spectral power in the higher 0.14 to 0.40Hz frequency bands), lower on-task RSA was linked to better performance. The authors suggested that, “reduced cardiac vagal tone during task execution helps establish an ergotopic [sympathetic-dominant] physiological condition, which contributes to optimizing mental functioning” (p. 115). Moreover, while our task did not require an overt response, prior work has indeed shown that greater DMN deactivation (here associated with lower RMSSD) associates with better task-performance, including improved memory (Daselaar, Prince, & Cabeza, 2004) or reduced prevalence of task-unrelated thoughts (McKiernan et al., 2006).

The analysis of BOLD correlates of RMSSD identified both the central and rostral cingulate gyrus bilaterally. The central part of the cingulate gyrus has been implicated in autonomic control. In particular, Critchley et al. (2003) found that ACC activity correlated with fluctuations in HRV during task performance. In addition, ACC-lesioned individuals did not show the typical task-related reduction in heart-rate variance shown by controls (and found in our current study). The RMSSD analysis also identified a very ventral part of vmPFC extending into medial orbitrofrontal cortex on the left and the subgenual ACC (sgACC). These regions have been linked to autonomic/emotional responses by Amodio and Frith (2006), and to sympathetic and parasympathetic activity in several studies (Nagai et al., 2004; Wong et al., 2007; Ziegler, Dahnke, Yeragani, & Br, 2009). Wager et al. (2009) reported that the magnitude of deactivation in the right orbitofrontal cortex (and putamen) was linked to higher heart rate (lower BBI) across individuals. We found a similar pattern for BBI though in a somewhat more rostral section (see Figure 4C), where increased heart rate was associated with greater activity. Thus, in this region, lower activity was linked to both lower heart rate and lower RMSSD.

As a whole however, the regions we identify for RMSSD overlap with the topological distribution of the Default Mode Network, a set of regions often associated with task-induced deactivation. This means that individuals who maintained higher vagal tone during task performance showed weaker task deactivation. Importantly, greater RMSSD *did not* correlate with stronger activation in any of the task-activated regions. Taken together this suggests that the link between deactivation and RMSSD is not associated with general attention, as in that case RMSSD should have also correlated with increased activation. Rather, less complex cardiovascular dynamics in response to task performance (i.e., reduced RMSSD) are specifically related to greater deactivation of the DMN.

It may be that individuals showing greater RMSSD during task performance are less engaged in task-related computations, which is accompanied by weaker task-induced deactivations in the DMN. As mentioned in the introduction, Wager et al. (2009) found that rapid fluctuations in the DMN correlate with autonomic state, which is consistent with a link to rapid shifts in attentional or homeostatic states. Apart from this, there is a separate literature linking the DMN to ANS function on short temporal scale. Using MEG, Park et al. 2014 showed that neural activity in the two main nodes of the DMN, the ventral ACC and right inferior parietal lobule, varied with the magnitude of the heart evoked response (HER) and predicted performance in a visual perception task. In addition, Babo-Rebelo et al. (2016) found that the proportion of self-focused thoughts during rumination covaried with HER amplitude and could be localized to the vicinity of the left precuneus. This suggests that the DMN mediates both physiological and cognitive functions.

We note that this account is somewhat independent from explanations of ACC involvement in modulation of sympathetic activity, as RMSSD loads predominantly on higher cardiac frequencies related to the parasympathetic system. With respect to sympathetic activity, it has been suggested (Critchley et al., 2003) that during normal function, ACC (dorsal part) controls the sympathetic system, and that reduced ACC activity is associated with lesser ANS control. Thayer et al. (2009) similarly suggested that disinhibition of prefrontal regions produces disinhibition in the central nucleus of the amygdala, producing a cascade of activations resulting in increased heart rate and decrease in HRV. Our findings are generally consistent with this as individuals with higher RMSSD showed weaker deactivation in these regions.

The areas identified in this analysis are more extensive than those documented by Beissner et al.‘s (2013) meta-analysis. For cognitive tasks, that analysis identified high-frequency HRV (parasympathetic) activity in the anterior insula, and in addition, sympathetic indices in a few other regions including vmPFC. That distribution is quite different from what we find. While it is difficult to relate any single set of findings (such as ours) to those of a meta analysis, perhaps the clearest difference is that the analysis method in our study may identify areas not be identified by typical reports. Typical analyses identify, by and large, areas where ANS activity tracks the task demands in a way that holds quite consistently at the group level. In contrast, the regions we identify are ones that can have a different feature these regions may show task-related activation/deactivation for some participants; those with the strongest ANS indices. This may be the main point of difference between our findings and those reported in Beissner et al.s meta-analysis.

#### 4.3.2 Correlation of other ANS measures with task-induced activation and deactivation

All ANS measures we examined identified brain systems for which ANS/BOLD correlations were either positive or negative. For the heart rate regressor (BBI) we found that faster heart rate correlated with task-induced activation in right fronto-parietal regions, which were strongly task-activated at the group level (compare Figure 4B and Figure 3). Faster heart rate was also linked to greater deactivation in right vmPFC (Figure 4C). Increases in heart rate are mainly caused by increased sympathetic activity, and are consequently related to a decrease in the vagal regulation (Hainsworth, 1995).

These findings for the BBI regressor are very consistent with several studies that have documented inverse correlations between vmPFC activity and arousal, either during task performance or the resting state. In a study of resting-state, Zeigler et al. (2009) found that faster heart periods were accompanied by lower activity only in the vmPFC (no area showed a positive correlation) Shmueli et al. (2007) found that fluctuations in HR tracked activity in several regions of the DMN (showing a negative correlation), mainly within ±6 sec of the measured BOLD response, but interpreted these as effects of physiological artifacts (but see Iacovella and Hasson, 2011, for a discussion on whether such correlations should be treated as physiological noise). Wong et al. (2007) found that engagement in a physical task produced changes in heart rate, which were negatively correlated with vmPFC activity. Furthermore, while the physical task used by Wong et al. deactivated both the vmPFC and the posterior cingulate cortex (PCC), which are both central DMN nodes, only vmPFC activity tracked heart rate, which is broadly consistent with our findings as the PCC was noticeably a region that did not show BOLD/ANS correlations for any of our autonomic measures. Nagai et al. (2004) examined correlations of skin conductance levels (SCL) with BOLD activity during a biofeedback task and found that increased SCL was associated with greater vmPFC deactivation. Taken together with those results, we suggest that the observed BBI-IID pattern of a positive ANS/BOLD correlation in task-active regions and a negative ANS/BOLD correlation in task-deactive regions is consistent with an attentional factor, and reflects a shift of the sympatho-vagal balance towards sympathetic activation and vagal deactivation.

Notably, the decrease of parasympathetic activity in task-deactive regions to an extent proportional to the degree of deactivation was also observed for respiration power. This ANS measure, which closely reflects the tidal volume, is an index of parasympathetic activity that has an impact, among other factors, on the respiratory sinus arrhythmia (Brown, Beightol, Koh, & Eckberg, 1985). Thus, the positive correlation found between the BOLD activity and the respiratory power in task-deactive regions (Figure 9) suggests that stronger deactivation occurred in subjects in whom a lower tidal volume was indicative of a weaker involvement of the parasympathetic control.

Interestingly, another ANS regressor – respiration frequency (fRESP) – also showed positive correlations for task-active regions and negative correlations for task-deactive regions, but for a different set of regions than identified by BBI. It is known that fRESP increases with task stress (Nilsen, Sand, Stovner, Leistad, & Westgaard, 2007) and that cognitive tasks including mental arithmetic increase respiration frequency (Wientjes, Grossman, & Gaillard, 1998). In our study, increased fRESP positively correlated with task-induced activation in visual cortices, and was negatively correlated with task-induced deactivation in temporal regions of the superior temporal plane as well as midline regions. The physiological mechanism behind this behavior may be again a decrease in the parasympathetic nervous activity: since higher breathing rates are usually associated with weaker respiratory sinus arrhythmia (Brown et al. 1985), the strong correlation between fRESP and the magnitude of task-induced (de)activation is likely related to the decrease in the vagal tone during successful responses to the task. The temporal regions in which deactivations correlated with fRESP are not generally linked to task related deactivation, but have been linked to modulation of auditory attention (Petkov et al., 2004), and their magnitude of activation follows the magnitude of pupil dilation during effortful listening (Zekveld, Heslenfeld, Johnsrude, Versfeld, & Kramer, 2014). Taken together with the positive correlation between fRESP and task-evoked activity in visual cortex, we suggest that BOLD/fRESP correlations may be linked to visual imagery processes that might be invoked during mental arithmetic. Amedi et al. (2005) showed that visual imagery is accompanied by activation of visual cortex and deactivation of auditory cortex, similar to the pattern we document. Amedi et al. also reported that increased activity in visual cortex correlated with decreased activity in auditory cortex, which is exactly the IID pattern found here. It is also possible that lateral temporal regions are more directly linked to the ANS. Duggento et al. (2016) reported a Granger-causality analysis between brain and cardiac activity and found that activity in several brain regions preceded fluctuations in the higher-frequency cardiac band, with 3 of these regions being lateral temporal ones (left MTG, right transverse temporal gyrus, right superior temporal pole).

As can be seen in Figure 2, there was no correlation between fRESP and BBI across participants (Pearsons *R* = –0.086), suggesting they load on different factors. This could explain why both were related to activation and deactivation but in different systems. It may be that the BBI measure loads on a more general attention-related factor, whereas the fRESP loads on cognitive components more strongly linked to working memory operations and imagery. Clearly such an account is speculative, but it suggests interesting directions for future research.

For cSE, we found that lower values, indicating less predictable cardiovascular variability, were linked to higher task-related activity in SFS bilaterally, central insula and central cingulate. The SFS areas showing these correlations were adjacent to those that showed task-related activation in the group-level analysis (the cluster on the right showed significant activation prior to FDR correction). The absence of any DMN region in these statistical maps, nor any area showing significant deactivation, suggests that cSE tracks BOLD activity in areas that show above-baseline activation for some participants, and may be generally related to attentional differences. The cSE reflects the complexity (unpredictability) of the portion of heart rate variability that is unrelated to respiration (Faes et al., 2015). Hence, the fact that the regions showing BOLD/cSE correlations exclude the DMN suggests that ANS indices accounting for vagally-mediated respiratory effects and respiration-unrelated (mainly sympathetic) effects load on different regions. This also provides further support to the role played by the respiration-related vagal ANS modulation documented by RMSSD (Figure 6) and pRESP (Figure 9).

The LF/HF ratio was relatively weakly correlated with all other measures. It identified only positive correlations with task-induced changes in two regions (left AG and vmPFC), which were task-deactive. Higher LF/HF values could be related to increased power in LF, perhaps due to sympathetic increase with stress (Bernardi et al., 2000), or alternatively, a slower breathing rate (thought in the current study, correlations with respiration frequency were moderate, Pearsons *R* = –0.37).

### 4.4 Implications for theories of individual-differences in deactivation

Certain populations show weaker-than-normal task-linked deactivation, particularly in the DMN. A good example is seen in research on schizophrenia, where several studies had reported a “failure to deactivate” (FTD) in schizophrenic populations (see Pomarol-Clotet et al., 2008). However, the interpretation of this finding has been unclear – deactivation in schizophrenia is *not* linked to task performance (Pomarol-Clotet et al., 2008) and for this reason, FTD in this population has been suggested to be generally related to the disease itself, perhaps indicating use of different computations, or less efficient cognitive performance. Similarly, landmark work in the study of autism (Kennedy, Redcay, & Courchesne, 2006) documented FTD throughout the DMN, which was interpreted in terms of abnormal internally directed processes at rest. However, later work on autism documented similar FTD patterns in behaviorally non-affected siblings of autistic individuals (Spencer et al., 2012), which suggests that FTD in autism may be related to a heritable feature that loads on a non-cognitive aspect related to deactivation. Chronic pain is associated with weaker deactivation in key nodes of the DMN (Baliki, Geha, Apkarian, & Chialvo, 2008), while accompanied by similar levels of task-based activation. Qin et al. (Qin, Hermans, van Marle, Luo, & Fernandez, 2009) reported that stress can impact the magnitude of deactivation in key DMN regions, with higher stress (as measured by cortisol concentration) accompanied by reduced deactivation. This has been interpreted in terms of a deficit in the typical of reallocation of resources induced by extrinsic tasks. The magnitude of deactivation reduces with maturation from childhood to early adulthood (Sun et al., 2013). Finally, reduced deactivation is also found in minimally conscious state (Crone et al., 2011).

As is evident from all these examples, reduced deactivation or FTD is found across the cognitive and clinical spectrum. However, the possibility that these failures are linked to, or are a result of abnormal autonomic responses under task demands has not been considered to date. Our findings suggest that a more complete understanding of failures to deactivate could be obtained by relating those to group or individual differences in maintenance of ANS activity. A parsimonious explanation is that these clinical states are associated with what amounts to a maladaptive maintenance of higher-complexity ANS states during task performance, which is reflected in reduced deactivation.

Finally, beyond accounting for inter-individual differences, such explanations may also account for interesting *intra*-individual differences in deactivation. A recent study (Meshulam & Malach, 2016) found that the progression of practice during a simple visual categorization task was accompanied by increased deactivation over time in the DMN. As mentioned by the authors, this may reflect task-related computations such as more successful silencing of competing information. Alternatively, as suggested by our findings, it may reflect lower cardiovascular involvement with practice. Relating such effects to ANS fluctuations within each participants scan is an interesting direction for future work.

### 4.5 Limitations and considerations for future work

Our study contains some limitations that could be annulled in future work. The small number of participants could result in lower power and subsequent misses relative to a larger cohort. However, we used robust regression so that the small number of participants would not necessarily increase the rate of false positives, as this type of regression method specifically down-weights univariate outliers. This regression procedure, while useful for allowing determination of the sign of the relationship and its statistical significance, does not allow precise estimation of effect sizes or confidence intervals (particularly for such a small sample), and future work using larger sample sizes would be necessary for these purposes (see *Supplementary Materials* for estimate of effective group size based on data from the current study).

There are also some technical limitations related to the acquisition of the physiological signals in our fMRI environment. The first is the utilization of the surrogate measure of HRV yielded by the plethysmogram (PPG). The BBI series that are the basis for mean-BBI, RMSSD, and LF/HF measures can differ when measured via ECG and PPG as a consequence of noise/artifacts and of the physiological variability of the transit time of the pressure wave from the heart to the peripheral location of PPG recording (Allen, 2007). As documented by Schafer et al. (Schafer & Vagedes, 2013), the extensive literature about the agreement between PPG- and ECG-based measures of HRV is not unequivocal. However, there does seem to be a consensus on the usability of PPG-derived measures of HRV in healthy subjects monitored in stable resting conditions (Selvaraj, Jaryal, Santhosh, Deepak, & Anand, 2008). It is unclear if this agreement maintains during physical or mental stress (Giardino, Lehrer, & Edelberg, 2002).

Another technical issue relates to the low sampling rate (50Hz) at which we acquired the PPG signal. This was determined by the scanner hardware, but is below that typically recommended for ECG-based HRV analysis (Task Force Of The European Society of Cardiology, 1996). Specifically, A low sampling rate reduces the accuracy in the detection of the fiducial points of cardiac events, which may cause changes in the subsequent derived autonomic indices. However, although sampling rate impacts the accuracy of HRV indices (Garcia-Gonzalez, Fernandez-Chimeno, & Ramos-Castro, 2004), traditional time, frequency and nonlinear indexes can be computed with a reasonable estimation error for low ECG sampling rates (Voss, Wessel, Sander, Malberg, & Dietz, 1996; Ziemssen, Gasch, & Ruediger, 2008), and even a sampling rate as low as 50 Hz can be used without irreparably degrading accuracy (Mahdiani, Jeyhani, Peltokangas, & Vehkaoja, 2015). Importantly, when we interpolated the PPG signal to a higher frequency, and recalculated RMSSD based on re-estimated peak locations, we found that the resulting RMSSD values were highly correlated with the original calculations suggesting that very similar BRAIN/ANS correlations would maintain under a higher sampling rate.

A separate issue relates to differentiating potential physiological effects on task-induced activation. In the current study, task performance had systematic impacts on autonomic measures, with two indices showing statistically significant effects, and three others statistically marginal ones. For this reason, we did not perform a procedure that partials out autonomic covariates from the BOLD signal. This procedure is typically referred to as ‘physiological noise correction’, and often used to account for ANS-induced fluctuations in BOLD resting state paradigms. However, as we have noted in prior work (Iacovella & Hasson, 2011), this procedure should not be automatically applied in task-related paradigms, particularly when it is known that the task produces ANS perturbations. In such cases, inserting ANS data as time-series regressors would result in regressing out meaningful activity patterns from exactly those regions involved in task-related computations. To evaluate this issue, we implemented such a correction on the BOLD time series collected in the current data (implementing a RETROICOR + RVT correction; see supplementary result) and re-calculated the group level activation patterns (analogous to those shown in Figure 3 in the main text). Implementing RETROICOR produced a clear pattern: it reduced the spatial extent of both activation and deactivation clusters as compared to basic findings, but at the same time it did not lead to identifying any new significant clusters.

While we capitalized on inter-individual differences in multiple aspects of heart rate variability, the current study does not address their causes. Explanations for inter-individual differences in HRV-related quantities are multi-factorial. They may be related to cognitive ability or strategy, physiology, or both (one factor underlying both physiology and cognition). Sampling participants in a way that provides sufficient data on such factors could allow understanding which factors underlie the relationship observed between HRV and brain activity. To this end, future work should consider controlling for such factors such as stress (Dishman et al., 2000) anxiety (Thayer, Friedman, & Borkovec, 1996), body mass (Karason et al., 1999), smoking habits (Hayano et al., 1990) and other factors known to vary with HRV.

Finally, the current study examined correlations between task-induced BOLD activity and IID in ANS indices during task performance. A comparable but separate question could probe for relations between BOLD activity and a ‘delta’ measure capturing the difference in ANS activity between task and rest. As opposed to this latter measure, the ANS measure we used likely reflects a combination of a tonic inter-individual factor related to overall function of the ANS system, as well as a phasic factor time-locked to task-induced ANS perturbation. Future work could focus solely on measures that capture task-induced perturbation to the ANS. In addition, our covert task did provide an indicator of participant’s effort or task performance. Future work could achieve this in different ways, including asking participants to press a key with each calculation step, recording the final number arrived at, at the end of all subtractions (to estimate the number of subtractions made) or employing of an independent behavioral task where participants produce verbal reports during the task.

### 4.6 Summary

We find that a meaningful proportion of IID in task induced deactivation correlate with IID in multiple autonomic constructs. Interestingly, different ANS measures correlated with task-induced deactivation (and to lesser extent, task-induced activation) in *different* brain systems. This shows that task-induced deactivation is multifaceted construct that can be better understood through its relation to different ANS measures. Finally, from the perspective of neuroimaging studies of ANS activity, our work shows the utility of deriving multiple ANS measures, as these load on different constructs related to task execution.

## Footnotes

1 Given the low sampling rate of the PPG, we interpolated that signal to 200hz, re-estimated the location of the fiducial peaks, and recalculated the RMSSD indices. The resulting set of values was very highly correlated (Pearsons *R* > .995) with RMSSD values estimated from the 50Hz series.

